# Atlantic Cod as an indicator of rapid climate change

**DOI:** 10.1101/2022.04.21.488963

**Authors:** Jae S. Choi

## Abstract

Atlantic cod (*Gadus morhua*) in the northwest Atlantic has become the textbook example of overfishing. However, this narrative has blinded us from the larger environmental context of this decline. Cod prefer cold and shallow habitats, environments that are also the most susceptible to rapid climate change. Cod habitat deterioration was evident well before their numerical decline, and as such, cod was a harbinger of rapid climate change. Recovery requires their habitat quality to first improve and stabilize. Calls for a cull of seals, the consensus actor causing the lack of recovery of cod, may therefore be unwise. Recovery will only occur once the variability associated with rapid climate change subsides and habitat quality improves. The timeline for this is of course unknown.

**One-Sentence Summary:** Was the decline of cod a harbinger of rapid climate change in the northwest Atlantic?

## Introduction

The collapse of Atlantic cod (*Gadus morhua*) in the North West Atlantic during the late-1980s has become the epitome of overfishing and its consequences [1–3]. Due to the strength of this conviction, a more precautionary and conservation-based approach towards natural resource exploitation evolved, with various area-based closures to fishing, a fishing moratorium upon cod, and eventually marine protected areas and reserves [4,5]. But the eagerly awaited return of cod did not manifest, even after three decades of effort. Almost right from the beginning, seals were noticed to be increasing in number and that they also can eat cod. Based upon this circumstantial evidence, they were considered the cause of the lack of return [6,7]. Of course, many other factors can influence cod mortality, including continued “subsistence” fishing and bycatch [8]; cannibalism, disease, and trophic interactions or cascades causing nonlinear predation release or density-dependent or “Allee”-type effects [9–14] effects. However, they represent variations on a theme: predation or consumer centric hypotheses. Seals just happen to be the predator of choice given the circumstances, leading to proposals for culls to help cod recover [11,15,16].

But what if this predation-centric mechanism is but a subplot, within a much larger story? Here we suggest this larger story is about cod habitat and how it’s environment has changed over the years. Indeed, in the years immediately following the collapse, anomalous temperature events were identified as a potential cause of the collapse [4,17,18]. This is because temperature preferenda are a profoundly important component of the habitat of all organisms. However, the variability in bottom temperatures were found to be enormous and an anomalous event of sufficient magnitude and sustained duration was not found to support the hypothesis ([5]; see Figures 1, 2). We suggest that rejecting this approach may have been premature. A more focused question needs to be asked, one that explicitly defines cod habitat and identifies its spatiotemporal evolution.

**Figure 1.**
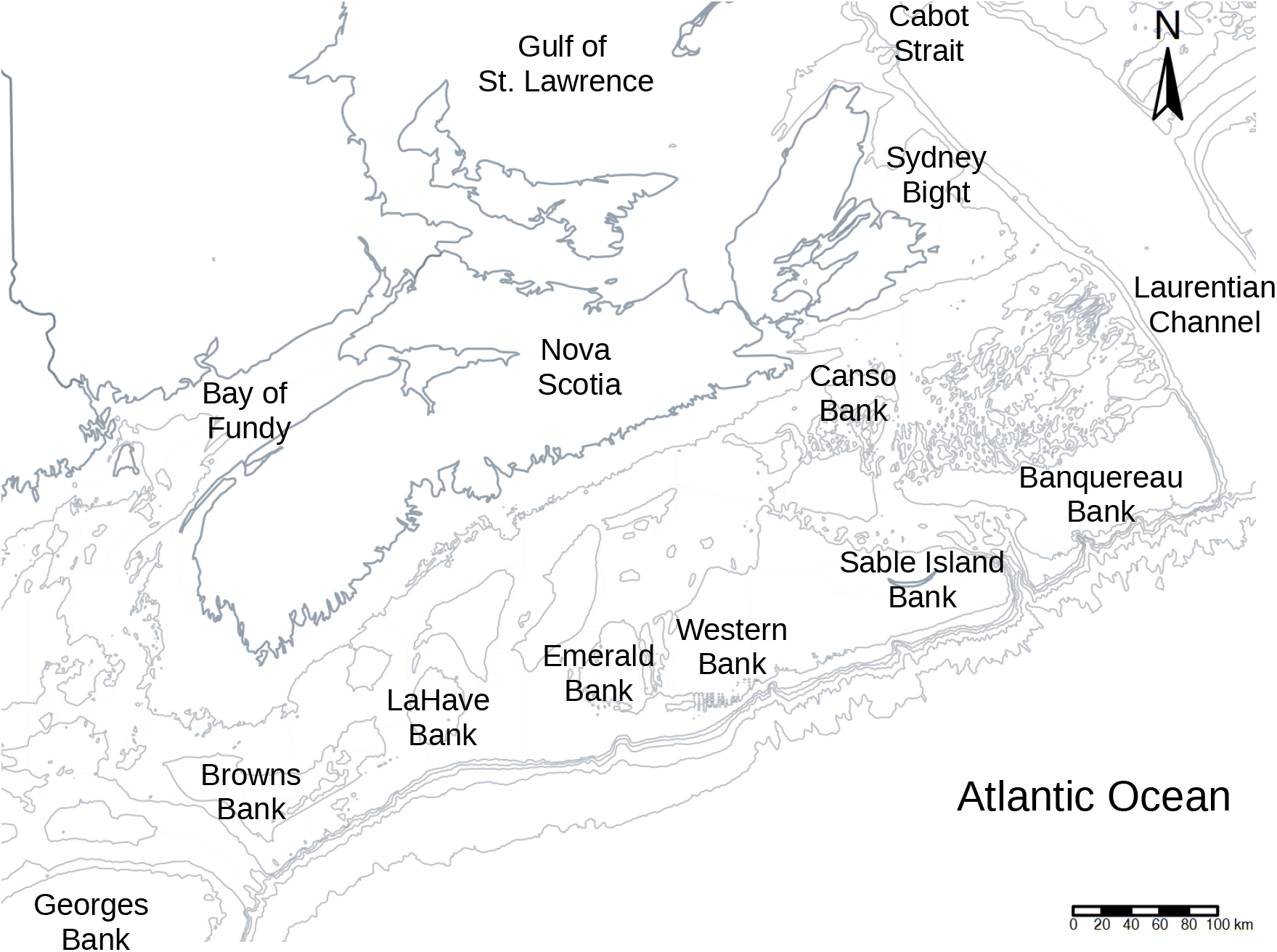
The area of interest in the Scotian Shelf of the northwest Atlantic Ocean (Canada, NAFO Div. 4VWX). Shown are isobaths and some of the major bathymetric features in the area. This area is at the confluence of the Gulf Stream from the south and south east along the shelf edge, Labrador Current and St. Lawrence outflow from the north and north east, as well as a nearshore Nova Scotia current, running from the northeast. It is hydro-dynamically very complex due to mixing of cold, low salinity water from the north with the warm saline water from the south.

**Figure 2.**
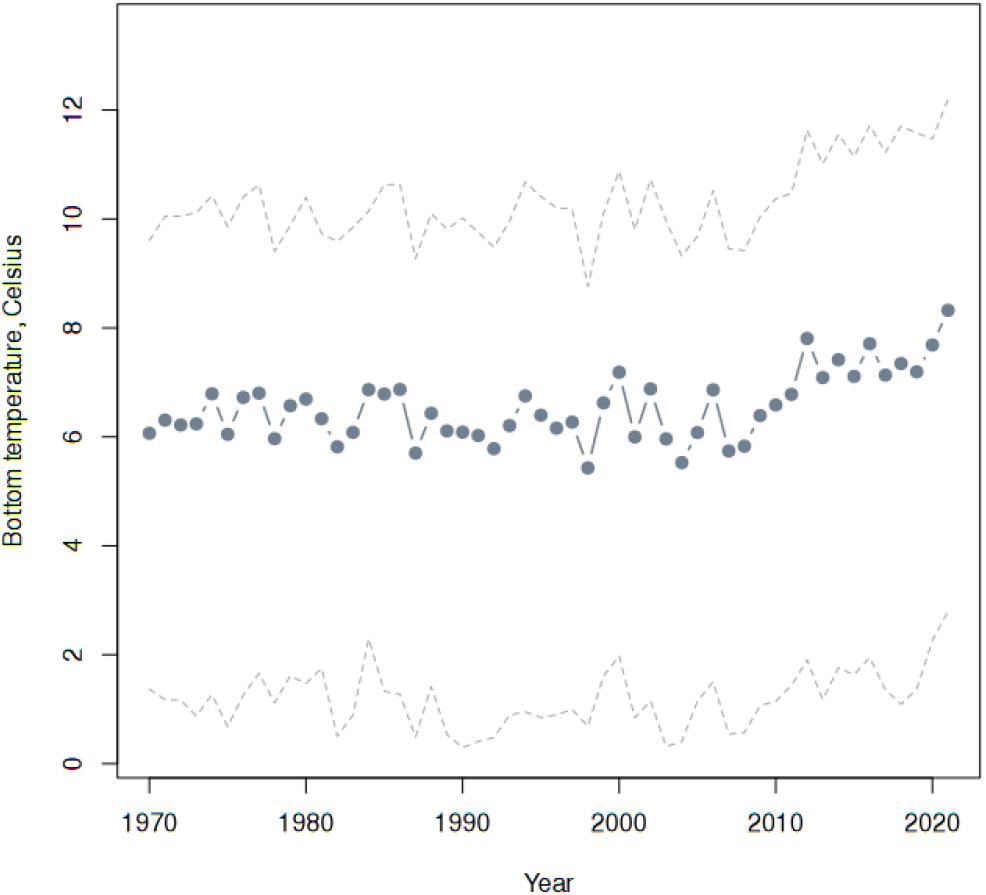
Posterior predicted bottom temperatures in °C on the Scotian Shelf with 95% Credible Intervals from depths of 50 m to 400 m, in the focal domain.

Fortunately, an explicit, probabilistic definition of habitat is readily computed as powerful analytical approaches have now become almost routine. Computational advances, most notably, the development of INLA [19] and the advent of spatiotemporal Conditionally Autoregressive (CAR) models that can borrow information from adjacent areas and times while operating in a Bayesian context [20–22] permit this definition. We use this approach to describe the spatiotemporal variations of cod habitat, focusing upon the Scotian Shelf of Atlantic Canada (NAFO Div. 4VWX; Figure 1).

## Materials and Methods

Specifically, we define “habitat” as a probability. When the probability of observing cod is greater than some threshold density, we can consider the space and time sampled to be “good habitat”, and those where it is less than this threshold to be “poor habitat”. Heuristically, we define this threshold to be the 0.05% empirical quantile of the non-zero densities of cod for a given type of sampling gear.

The binary categorization of each sampling event at an areal unit s=( 1,…, a ) and time unit t=( 1,…, b ), Y_st_, is modeled as a *Bernoulli* process with a *logit* link function and conditional upon covariates and autocorrelated random effects in space, time and spacetime:

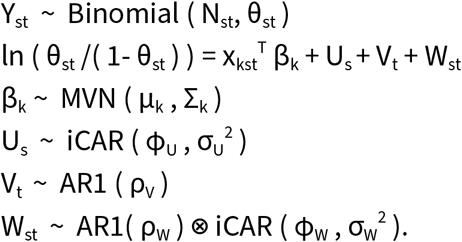

Here, θ_st_ the vector of probabilities of success in each trial in each space and time unit and N_st_ the number of trials; the 3D matrix of covariates x_kst_^⊤^ and covariate coefficients parameters β_k_ with a multivariate normal prior with mean μ_k_ and diagonal variance matrix ∑_k_; the spatial autocorrelation U_s_ follows the iCAR (also known as a convolution model or the Besag-York-Mollié or “BYM” model in the epidemiological literature, when data are Poisson distributed) with φ_U_ the proportion of the iCAR variance, σ_U_^2^, attributable to spatial processes relative to nonspatial processes; the temporal error components V_t_ are assumed to follow a lag-1 autocorrelation process (AR1) with parameter ρ_V_; and finally, the spatiotemporal errors W_st_ are modeled as a Kronecker product (⊗) of temporal AR1 and spatial iCAR processes, where time is nested in space. This model form is known as an “inseparable” space-time model.

For these analyses, we drew upon two high-quality bottom trawl-based research surveys conducted on the Scotian Shelf of Canada, in the Northwest Atlantic Ocean. The first is an annual multispecies survey conducted by the Department of Fisheries and Oceans (DFO) from 1970 to the present (“ground fish survey”). Sampling stations were randomly allocated in depth-related strata with sample tows of approximately 1.82 to 2.73 km in length on a variety of different gears, mostly the Western IIA net. Vessel speeds ranged from 4.5 to 6.5 km/hr. Only tow-distances were availability which was used to approximate areal density using a nominal wing spread of 12.5 m.

The second data series is a survey conducted through a joint collaboration between the snow crab industry in the region and DFO, from 1999 to the present (“snow crab survey”). The survey is funded by a proactive industry as a forward-looking investment towards long-term sustainable fishing, an exemplary model of the precautionary approach to fishing. A modified Bigouden Nephrops net designed to dig into sediments was used with sample tows of approximately 0.3 km in length and a target vessel speed of 3.7 km/h. Actual bottom contact and net configuration were monitored to provide explicit areal density estimates. The wing spreads ranged from 12 to 14 m, depending upon the substrate encountered.

Amongst the *k* covariates in this study were: bottom temperatures, depth, gear type, season, and a global intercept term as the overall average habitat quality. The Western IIA was used as the reference gear, the longest running survey gear.

As multiple surveys with different sampling designs were used, a hybrid, Voronoi triangulation and tessellation-based solution was used. The algorithm maintains an approximately constant station density within each areal unit, a target of 30 sampling events within each areal unit and a final total number of areal units of less than 2000 units. This was a trade-off between information density and computational stability. The tesselation-based approach had the advantage of more informative areal units capable of resolving temporal trends and environmental gradients. This was critical as environmental variability has been significant in the area/time.

All analyses were conducted with R and R-INLA with all code available online (https://github.com/jae0/aegis.survey/blob/master/inst/scripts/10b_cod_carstm_tesselation.R). Note that we utilize the “bym2” parametrization of the iCAR for stability and interpretability of parameter estimates [21]. Penalized Complexity (PC) priors that shrink towards a null (uninformative base model) were used for all random effects [23]. For the AR1 process, we used the “cor0” PC prior which has a base model of 0 (i.e., no correlation). For the spatial process (“bym2”) we use the PC prior (rho=0.5, alpha=0.5). Seasonality was discretized into 10 units and a cyclic “rw2” process was used. For all other covariates, a second order random walk process (“rw2”) was used as a smoother using 11 quantiles as discretization points.

## Results and Discussions

The posterior parameter estimates were biologically informative and meaningful (Table 1). Overall, there was a 65.4% probability (intercept term) of observing cod habitat in the spatial domain, from 1970 to 2021. In order of importance (variance components), the factors that most influenced habitat quality for cod were: persistent random spatial effects, spatiotemporal random effects, depth, sampling gear, interannual variability, temperature and seasonal cycles. The overall temporal (inter-annual) autocorrelation across time (ρ) was high (0.945) and marginally smaller when resolved to areal units (0.909) attributable to local processes. The relative importance of neighborhood effects (φ) was high when time was not resolved (0.989) and when time was resolved (0.982).

**Table 1.** Posterior parameter mean and 95% Credible Intervals from the habitat model. σ is the posterior estimate of the standard deviation estimated for each process. ρ is the posterior temporal autocorrelation parameter and φ is the posterior proportion of the spatial and space-time variability associated with neighbourhood effects.

| Parameter estimates | Mean | Quantile 0.025 | Quantile 0.975 |
| --- | --- | --- | --- |
| Intercept | 0.654 | 0.274 | 0.914 |
| $\sigma$ time | 0.715 | 0.483 | 1.048 |
| $\sigma$ cyclic | 0.374 | 0.176 | 0.641 |
| $\sigma$ gear | 0.740 | 0.357 | 1.431 |
| $\sigma$ bottom temperature | 0.406 | 0.218 | 0.667 |
| $\sigma$ depth | 0.793 | 0.377 | 1.540 |
| $\sigma$ space | 1.383 | 1.235 | 1.521 |
| $\sigma$ space_time | 1.188 | 1.072 | 1.332 |
| $\rho$ time | 0.945 | 0.887 | 0.980 |
| $\rho$ space_time | 0.909 | 0.877 | 0.940 |
| $\phi$ space | 0.989 | 0.970 | 0.998 |
| $\phi$ space_time | 0.982 | 0.952 | 0.997 |

By accounting for these factors, albeit in the above admittedly simplistic model, we can begin to disentangle the *realized* thermal preferences of cod from other factors (Figure 3). In our focal spatial domain, the highest probabilities (0.65) of observing cod are from 4 to 5 °C and diminishing to a probability of less than 0.5 when temperatures are colder than 2 °C and warmer than 7 °C. But cod habitat is not defined by temperature alone. Depth is a well-known proxy for pressure, light, turbulence, substrate complexity, and overall variability of any number of other environmental factors. We find that cod has a preference for shallower environments, with the probability of finding cod dropping off to less than 0.5 beyond a depth of 150 m (Figure 3).

**Figure 3.**
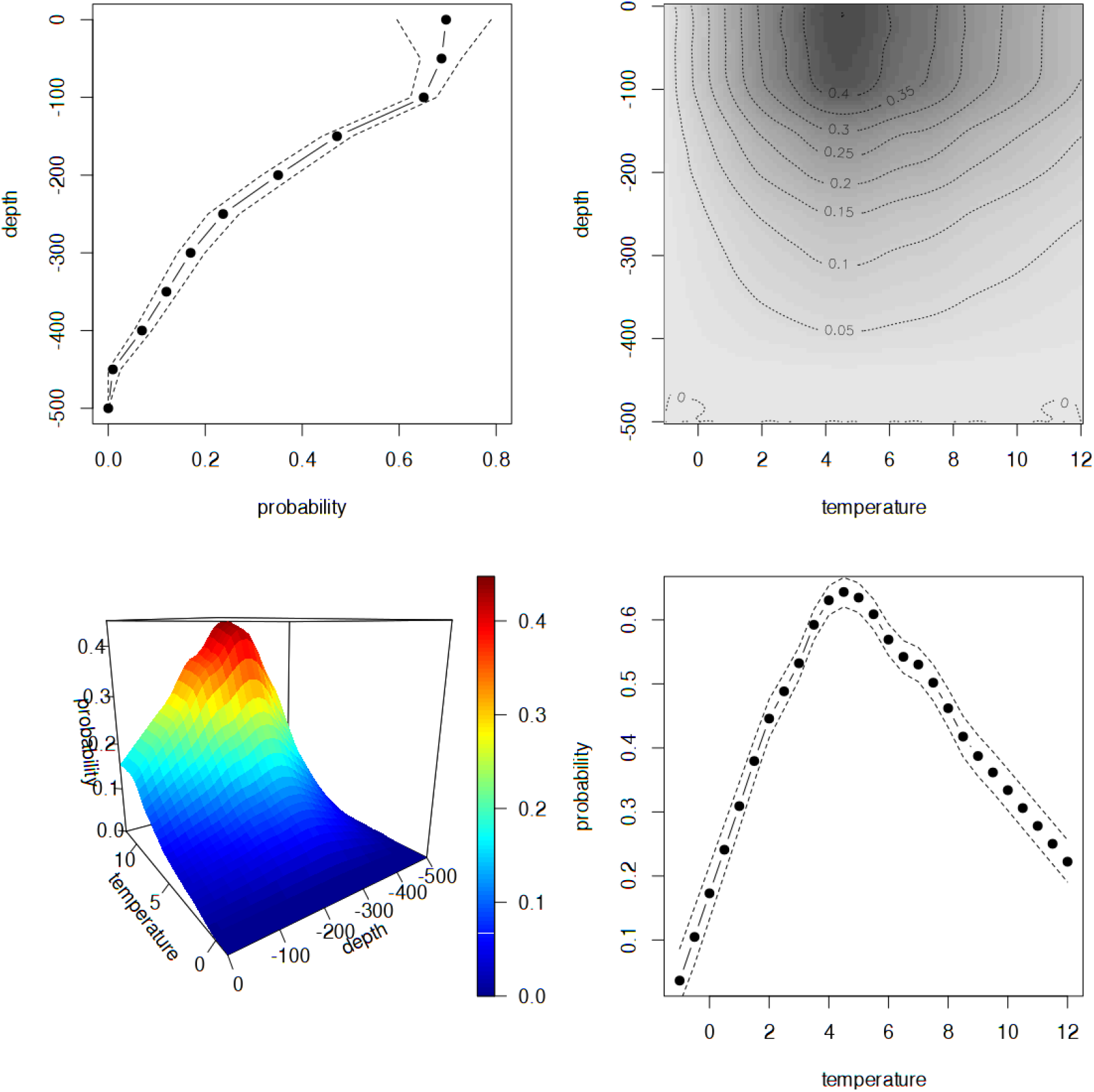
Posterior predicted relationship of the probability of a location being good cod habitat as a function of bottom temperature (°C; bottom right), and depth (m; top left). Stippled lines are 95% Credible Intervals. Top right is a contour of the probability of habitat and the bottom left is a 3D visualization of this pattern. Note peak probability at depths less than 100 m and at 4 °C.

Clearly, cod in the Scotian Shelf region have a preference for shallow and cooler habitats (less than 80 m and between 4 and 5 °C). This narrow habitat preference also renders cod more susceptible to rapid climate change. Indeed, in the spatial domain of our study, the habitat quality for cod has consistently declined since a peak in the mid-1980s (Figure 4). The overall average probability of finding cod habitat fell below 0.5 by the mid-1990s and currently remains at 0.3. The timing of the decline in numerical abundance in the early 1990s coincides with the degradation of habitat. It is a systemic degradation of cod habitat probabilities.

**Figure 4.**
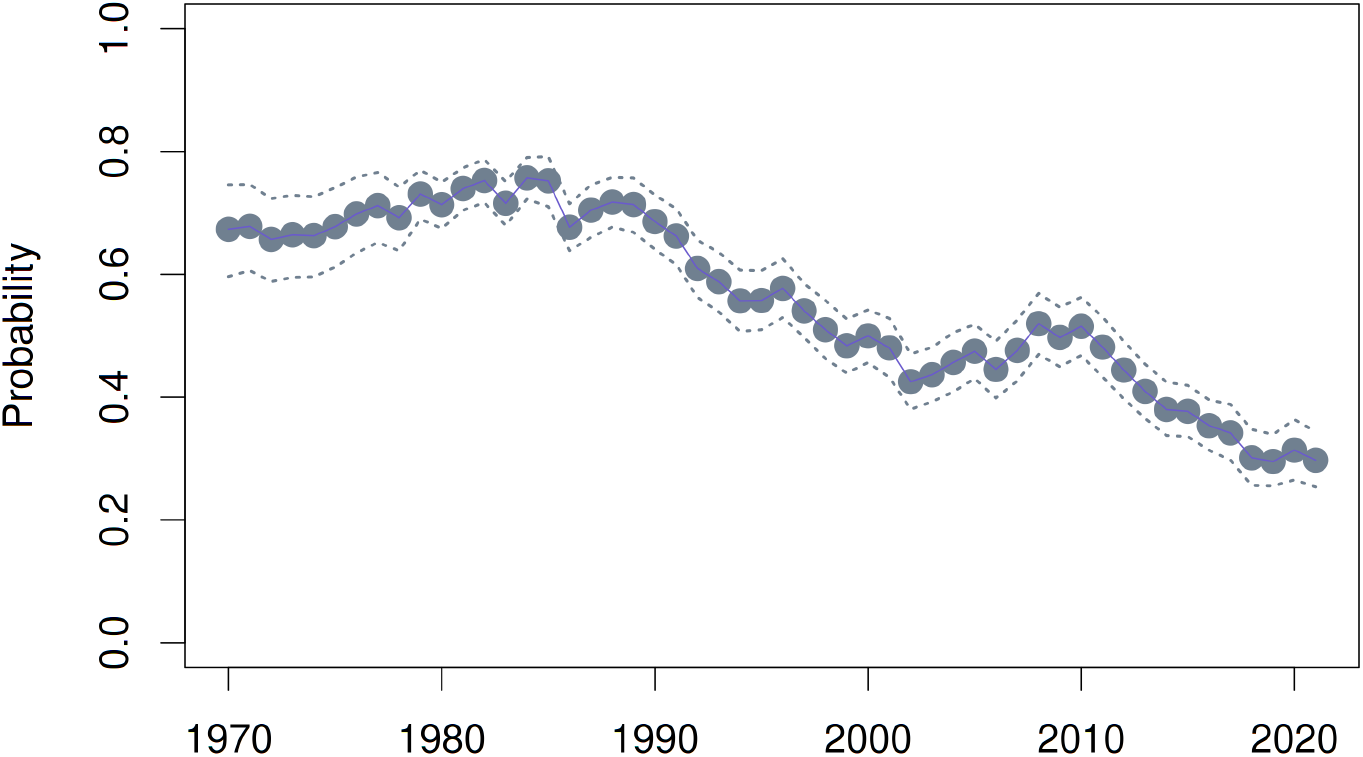
Posterior predicted mean cod habitat probability on the Scotian Shelf (Summer Strata, excluding Georges Bank) as a function of time. Stippled lines are 95% Credible Intervals.

Decomposing the effects of depth, the locations where depth is less than 80 m (Figure 5) identify potential depth-related habitat. Similarly an application of the bottom temperature predictions and depth variations as determinants of habitat (Figure 6) identify their interplay as an important contributor to the decline in average habitat quality (Figure 4). The pure spatial effect (Figure 7) identifies persistently high habitat probability locations, that is, core habitat that have been most robust to habitat degradation. However, throughout the domain, the presence of optimal habitat for cod declined not only due to warming (above 8°C) but also cooling (below 2 °C) in other areas (Figure 6). That is, overall climatic variability jointly operating with depth to create suboptimal habitat. Where optimal temperatures did occur, they often occurred in areas of greater depth (> 150 m; Figure 5) which did not serve to improve cod habitat.

**Figure 5.**
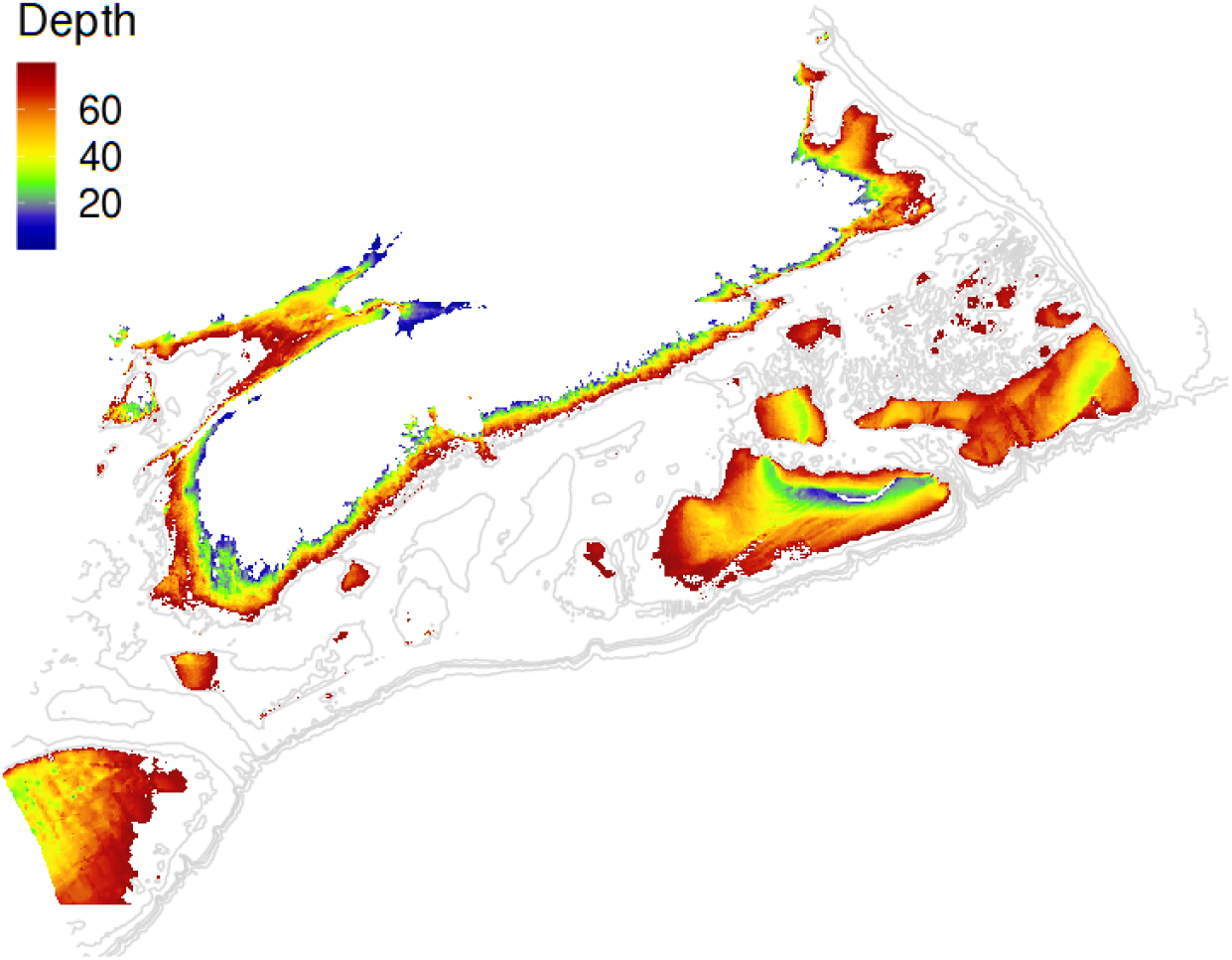
Locations with depths between 10 and 80 m, depths with the highest probability of being cod habitat.

**Figure 6.**
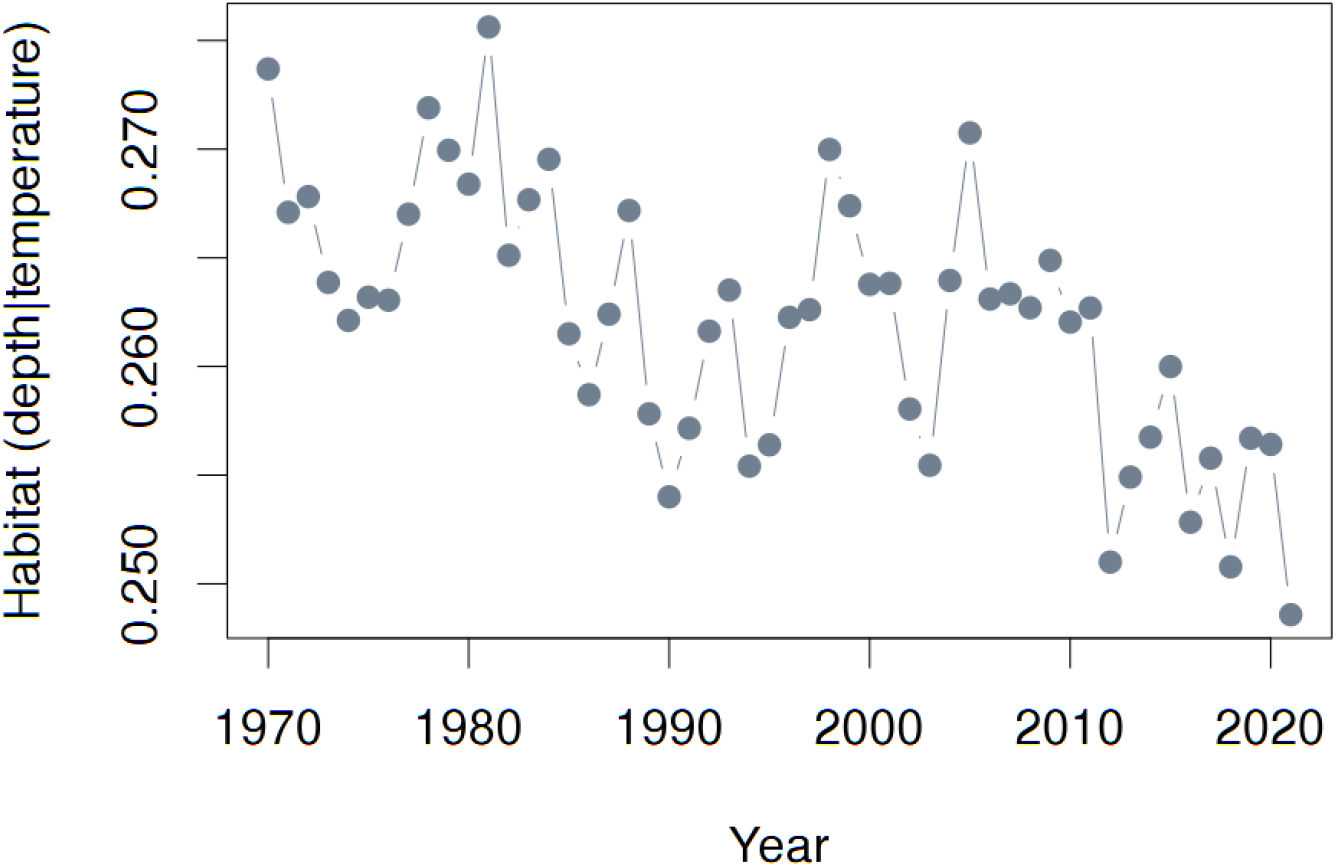
Mean habitat probability based upon depth and bottom temperature. This extraction is for the “Summer Strata”, that excludes Georges Bank. Note the extreme low in 1990, near the period of rapid numerical decline.

**Figure 7.**
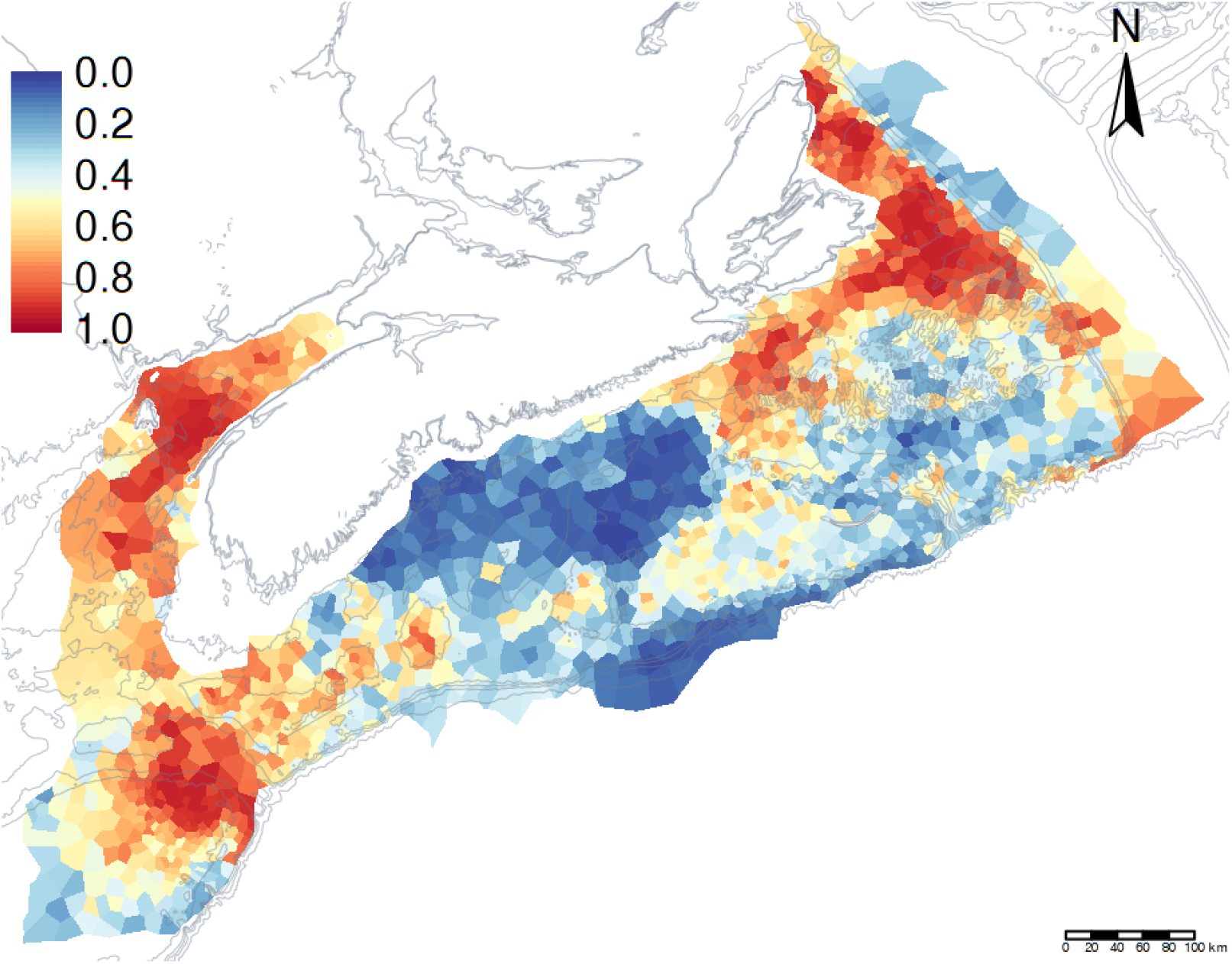
Posterior marginal spatial effects on cod habitat probability; that is, persistent spatial patterns of habitat or “core” habitat areas.

Variability in detection probability of habitat was also evident with gear type (Figure 8) with a large increase in the probability of detecting cod habitat from the Western IIA (historically, the primary survey gear) relative to the US 4 seam 3 bridle survey trawl net (the current standard). The Nephrops net used in the snow crab survey was intermediate in ability to capture cod.

**Figure 8.**
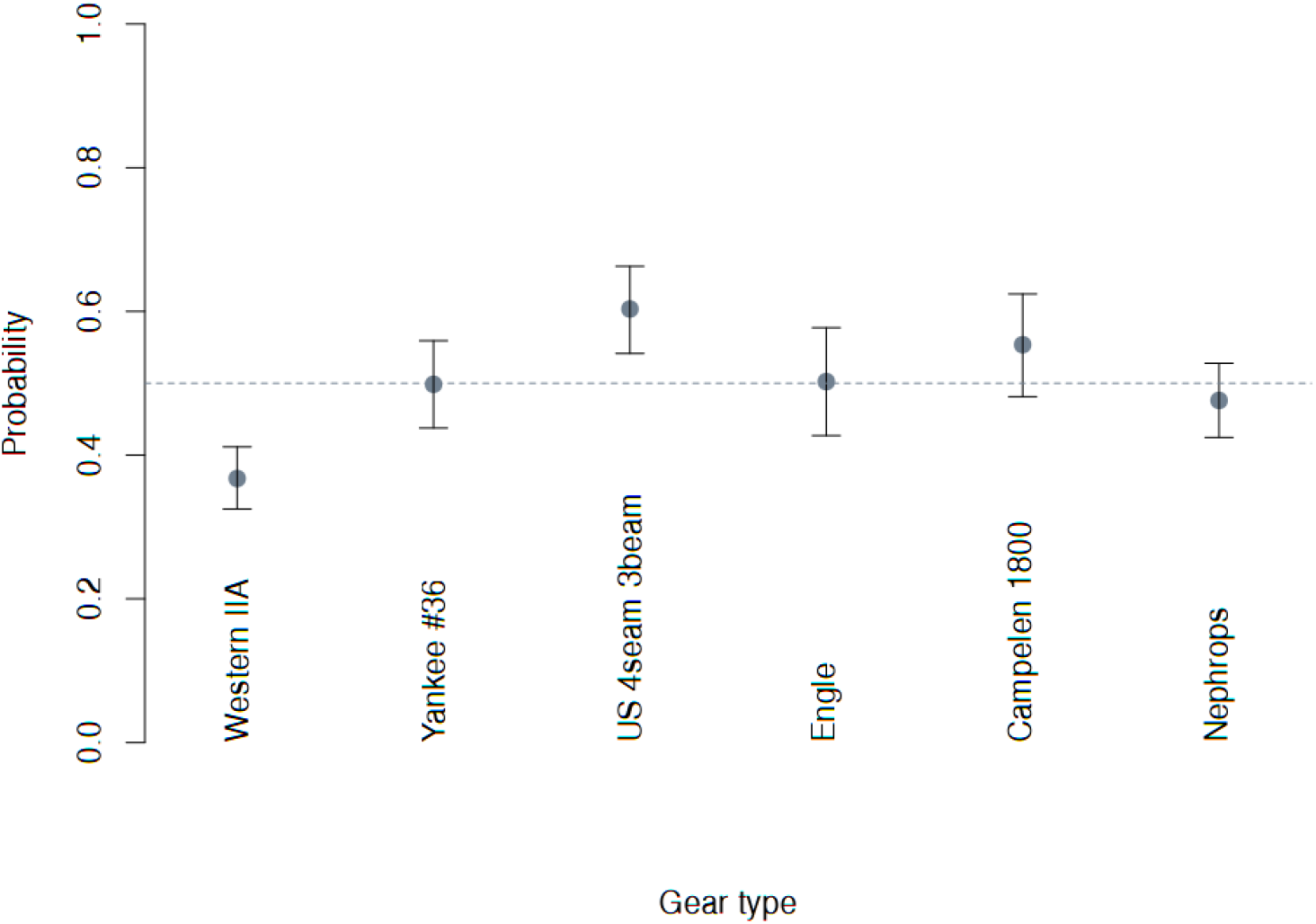
The posterior marginal effect of sampling gear upon the probability of finding cod habitat, independent of all other factors. Note that most have overlapping 95% Credible Intervals, though some strong differences are found between the Western IIA (most frequently used gear in the past) and the US 4 seam 3 bridle survey trawl net, the current standard.

In summary, from a habitat perspective, fishing and seal predation along with any other mortality factor, only exacerbated the decline of cod as the preconditions necessary and sufficient for their decline were already written: spatial and temporal variability of bottom temperatures. With such habitat decline, correlated markers of a population in distress followed: reduced population size, stunted growth, early maturation, disease/parasite prevalence, declines in physiological condition [5], and most importantly, declines in the capacity to be resilient and resistant to these perturbations [24]. Most of these latter issues have been historically blamed squarely upon fishing. There is a need to recontextualize these processes in the context of habitat change and stress. It remains to be seen if similar processes are responsible for the collapse and lack of return of cod in other areas of the northwest Atlantic. However, the large scaled nature of these climatic processes suggests that they will likely play an important role elsewhere.

## Conclusions

The outcome of the collapse was a greater awareness of the ecological frailty of even historically very dominant fisheries and the general adoption of more precautionary approaches to fishery exploitation. These are all well and good. However, the real issue seems actually to have been a much more significant large-scale process: rapid climate change causing spatiotemporal variability in habitat. The premature presumption that only predator-prey interactions control cod, whether it be human, seal or other, has prevented the wider lens of rapid climate change and its influence upon cod habitat [17] from being perceived. Indeed, the demise of cod may be better contextualized as having been an early indicator of significant environmental change. As early as the 1990s cod were suffering from the reality of significant oceanic climate variability (Figures 4, 6), which we were, unfortunately, unable to appreciate in time.

If this larger narrative is correct, reversion to a cod-dominated system is, of course, possible. However, first and foremost, habitat conditions must revert to much higher probability levels. This must also last for a minimum of five years, the time to reach maturity, in order for strong year classes to form. Some minor reversion in habitat quality did occur prior to 2009 (Figure 4), possibly associated with extended La Nina events. However, this did not last long enough to have a significant biological effect upon cod, as they were quickly followed by a succession of extended El Nino events and conditions more extreme than that associated with rapid declines in 1990 (the years 2012, 2016, 2018 and 2021 in Figure 6).

Of course, predation by humans, seals and any other mortality factor also need to cooperate; any conservation and marine spatial planning measures for cod must focus upon such shallower and intermediate temperature core habitat environments. Irrespective, it must be understood that the underlying causes of these habitat changes are occurring on large time and spatial scales, via processes over which we have limited immediate and direct control. Culling seals or any other perceived predator will likely have no influence upon recovery if there is no suitable home for them to occupy. For now, we must accept that a systemic, environmentally driven hysteresis has occurred [25].

## Data and materials availability

All data are held in databases in Fisheries and Oceans Canada. All analytical code are available at https://github.com/jae0/aegis.surveys/.

The perspectives outlined here are those of the author and not necessarily that of funders nor Fisheries and Oceans Canada.

